# Reference-guided comparative genomics of seven Indonesian rice cultivars identifies conserved gene space and trait-associated sequence candidates

**DOI:** 10.64898/2026.08.26.747264

**Authors:** Yekti Asih Purwestri, Adhityo Wicaksono, Siti Nurbaiti, Nada Tazkia Purba, Dewi Retnaningati, Ratih Restiani, Nirma Kumalasari, Tri Rini Nuringtyas, Valentina Dwi Suci Handayani

**Affiliations:** Department of Tropical Biology, Faculty of Biology, Universitas Gadjah Mada, Jl. Teknika Selatan, Sekip Utara, Sleman, Yogyakarta 55281, Indonesia; Research Center for Biotechnology, Universitas Gadjah Mada, Jl. Teknika Utara, Sleman 55281, Yogyakarta, Indonesia; Biotechnology Study Program, The Graduate School, Universitas Gadjah Mada. Jl. Teknika Utara, Sleman 55281, Yogyakarta, Indonesia; Department of Biology Education, Teacher Training and Education Faculty, Universitas Borneo Tarakan, Jl. Amal Lama No. 1, Tarakan 77115, Indonesia; Department of Agronomy, Faculty of Agriculture, Universitas Gadjah Mada, Jl. Flora, Bulaksumur, Sleman 55281, Yogyakarta, Indonesia

**Keywords:** Indonesian rice germplasm, PacBio HiFi, reference-guided consensus genome, comparative genomics, orthogroups, candidate-gene analysis

## Abstract

Indonesian rice cultivars represent valuable genetic resources, yet many remain poorly characterized at the genomic level. Here, we generated 95.40 Gb of PacBio HiFi sequence data from seven Indonesian rice cultivars and constructed cultivar-specific consensus genomes using the telomere-to-telomere Nipponbare reference AGIS1.0. Sequencing coverage ranged from 27.92× to 41.58×, and the resulting consensus genomes spanned 387.93-390.54 Mb, with BUSCO completeness of approximately 98.3-98.5%. OrthoFinder assigned 99.1% of predicted proteins to 40,737 orthogroups, including 27,514 core orthogroups represented across all seven cultivars, indicating a highly conserved predicted gene space within the reference-guided framework. Targeted analysis recovered 278 of 280 cultivar-by-locus combinations representing 40 genes or gene family entries associated with grain pigmentation, nitrogen and amino-acid metabolism, and starch properties. Comparative predicted protein analysis prioritized *ANS1*, *SBE2b*, *SSIIa/ALK*, *Wx/GBSSI*, *OsAAP6/qPC1*, and *SSI* as candidates for further investigation. Among 269 completed AGIS1.0-anchored promoter comparisons, 159 passed quality-control criteria, whereas 110 were flagged for gene-model, boundary, synteny, or structural concerns. Notably, these flagged comparisons accounted for more than 90% of the alignment-derived sequence variation, emphasizing the importance of rigorous quality control when interpreting apparent promoter divergence. Collectively, these reference-guided genomic resources provide a standardized framework for investigating sequence variation in Indonesian rice germplasm and prioritize testable coding and regulatory candidates for functional validation and future genomics-assisted crop improvement.

## Introduction

Rice is central to food security in Indonesia, but its importance extends beyond production volume. Local cultivars and landraces retain genetic variation shaped by geography, cultivation environment, farmer selection, food traditions, and cultural preferences. Indonesian rice germplasm encompasses both indica and tropical japonica lineages and can show substantial diversity even among accessions collected within a relatively restricted region (Thomson et al., 2009). Such locally maintained variation may contain useful alleles for environmental adaptation, grain quality, nutritional composition, and crop improvement that are poorly represented in intensively selected elite varieties. Large-scale rice genomics has demonstrated extensive population structure and millions of sequence variants across Asian cultivated rice, but locally important Indonesian cultivars remain comparatively underrepresented in genome-scale resources (Wang et al., 2018). Genomic characterization of these materials is therefore important both for conserving Indonesian rice diversity and for enabling its informed use in future breeding and functional studies.

The present study examined seven Indonesian rice cultivars: Inbrida Pari Gogo 12/Inpago (IN), Cempo Ireng (CI), Gogo Jak (GJ), Kisol Manggarai (KM), Merah Pari Eja (MPE), Boawae Seratus Malam (BSM), and Putih Payo (PP). The panel represents contrasting cultivation backgrounds, grain pigmentation, and previously reported physiological responses. CI is an Indonesian black-rice cultivar for which genomic, metabolomic, ionomic, transformation, and genome-editing resources have begun to be established (Sedeek et al., 2023). BSM, GJ, and KM are local cultivars from East Nusa Tenggara that have shown distinct physiological and transcriptional responses to drought, with BSM exhibiting particularly strong drought-tolerance characteristics in previous experiments (Salsinha et al., 2022). MPE is a pigmented red-rice cultivar previously identified as drought-tolerant, whereas the non-pigmented PP was highly drought-sensitive under the same screening conditions (Sebastian et al., 2022). Together with the accession recorded as Inpago, these cultivars provide a biologically diverse comparative panel, although they were not sampled to represent the complete population structure of Indonesian rice.

Whole-genome sequencing (WGS) provides a more comprehensive basis for cultivar comparison than morphological traits or a restricted set of molecular markers because it enables variation to be examined across coding regions, regulatory intervals, repetitive sequences, and the broader predicted gene space. PacBio circular-consensus sequencing produces HiFi reads that combine long-read continuity with high single-read accuracy, supporting reliable genome reconstruction and sequence-variant detection (Wenger et al., 2019). In this study, PacBio HiFi data were used to construct cultivar-specific consensus genomes against AGIS1.0, a telomere-to-telomere assembly of the Nipponbare rice reference in which all 12 centromeres and 24 telomeres were resolved (Shang et al., 2023). This reference-guided strategy provides coordinate-compatible genomic resources suitable for standardized annotation and locus-level comparison. However, because the resulting genomes retain the structure of AGIS1.0, they should be interpreted as cultivar-specific representations of reference-relative sequence variation rather than independent de novo assemblies or a comprehensive Indonesian rice pan-genome.

Candidate genes were selected from three trait-related biological groups with direct relevance to grain properties. The first comprised amino-acid transport, storage, and nitrogen metabolism, including amino-acid permeases, the GS/GOGAT system, *ASN1*, and *LKR/SDH*. These processes determine how assimilated nitrogen is converted, transported, remobilized, and ultimately allocated to developing grain; notably, OsAAP6 has been functionally associated with amino-acid distribution and grain protein content (Peng et al., 2014). The second group included anthocyanin-pathway genes such as *CHS*, *CHI*, *DFR*, and *ANS*, together with the carotenoid-pathway genes *PSY*, *LCYB*, and *LCYE*. Pigmentation was included because genetic changes affecting anthocyanin biosynthesis and regulation can produce major differences in rice grain colour and bioactive composition (Oikawa et al., 2015). The third group comprised starch-biosynthetic genes, including *SBE1*, *SBE2b*, *Wx/GBSSI*, and members of the SSI, SSII, and SSIII families. Natural variation in *Wx* and ALK/SSIIa can alter amylose structure, gelatinization temperature, mouthfeel, and other cooking and eating properties (Gao et al., 2003; Zhang et al., 2019). These loci were therefore treated as biologically informed candidates rather than as genes formally associated with phenotypes in the seven cultivars.

Accordingly, this study aimed to establish and comparatively characterize reference-guided genomic resources for seven Indonesian rice cultivars. We first evaluated the sequencing and consensus-genome datasets through mapping statistics, gene-space completeness, repeat profiling, structural and functional annotation, BUSCO-based phylogenomics, and orthogroup inference. We then examined the representation of conserved, semi-core, accessory, private, and reference-only predicted orthogroups and performed targeted recovery of genes involved in pigmentation, amino-acid and nitrogen metabolism, and starch properties. Candidate loci were further evaluated for predicted protein-sequence differences, annotation-quality concerns, and proximal promoter variation relative to AGIS1.0. The study was designed to identify conserved genomic features and prioritize cultivar-dependent coding, promoter, and gene-model differences for subsequent validation, and not to establish causal alleles or definitive gene presence-absence variation. The resulting resource provides a foundation for read-backed locus validation, transcriptomic and biochemical investigation, conservation of Indonesian rice germplasm, and future genomics-assisted improvement of locally important cultivars.

## Materials and Methods

### Plant materials, extraction, and sequencing

Seven Indonesian rice cultivars were examined: Inbrida Pari Gogo 12/Inpago (IN) from Indonesian Center for Rice Research (BBPadi), West Java; Cempo Ireng (CI) from Sleman Yogyakarta; Gogo Jak (GJ) from Jak Village, East Manggarai, East Nusa Tenggara (NTT); Kisol Manggarai (KM) from Kisol District, East Manggarai, NTT; Merah Pari Eja (MPE) from Tagari Village, Bulusu, North Toraja, South Sulawesi; Boawae Seratus Malam (BSM) from Nageoga Village, Boawae, Nagekeo, NTT; and Putih Payo (PP) from Lempur, Kerinci, Jambi. All obtained plant seeds were cultivated in the greenhouse of Universitas Gadjah Mada (UGM). Genomic DNA was extracted from mature leaf tissue collected from one plant per cultivar using the Quick-DNA™ Plant/Seed Miniprep Kit (Zymo Research, USA) according to the manufacturer’s protocol. DNA purity was assessed from the A260/A280 and A260/A230 ratios using a MaestroNano Pro microvolume spectrophotometer (MaestroGen, Taiwan). For WGS, DNA library preparation for Pacific Biosciences (PacBio) was used, with a Qubit fluorometer, and an integrity check pipeline was run by the service provider Integrated Genome Factory (IGF), Indonesia. The sequencing device was PacBio Revio, which generates long reads. The SMRT Link software for the PacBio HiFi sequencing device ran with smrttools v25.2.0.266456 (core v6.5.14) and basecaller v13.0.0.

### PacBio HiFi read mapping, variant calling, and reference-guided consensus generation

PacBio HiFi reads were delivered in BAM format and converted to FASTQ using pbtk v3.5.0. Sequence yield, read count, mean read length, and read N50 were calculated using SeqKit v2.13.0 (Shen et al. 2016). Reads were aligned to the AGIS1.0 Nipponbare reference genome (RefSeq accession GCF_034140825.1; Shang et al. 2023) using pbmm2 v26.1.99. Alignments were coordinate-sorted and indexed using SAMtools v1.22 (Danecek et al. 2021). Mapping statistics, including mapped-read proportion, mean mapping quality, and mean depth, were obtained using Qualimap v2.3 (Okonechnikov et al. 2015). Small variants were called using BCFtools v1.22. Genotype likelihoods were generated using bcftools mpileup, followed by genotype calling using bcftools call with consensus calling mode. Cultivar consensus sequences were generated using bcftools consensus. The resulting sequences were therefore treated as unphased, reference-guided cultivar consensus sequences rather than independent genome assemblies.

### Repeat identification, genome masking, and gene prediction

Repetitive sequences were modelled using RepeatModeler v2.0.5 (Flynn et al. 2020). A separate de novo repeat library was generated for each consensus sequence. The resulting repeat library was used with RepeatMasker v4.1.5 (Smit et al. 2013-2015) to soft-mask the reference-guided consensus sequences before gene prediction. Protein-coding gene models were predicted from the soft-masked consensus sequences using AUGUSTUS v3.5.0 (Stanke et al. 2008; Hoff and Stanke 2019). AUGUSTUS parameters for *Oryza sativa* were trained using the AGIS1.0 gene file format (GFF) annotation that is available in the NCBI Genome. The AGIS1.0 GFF annotation was used to establish the gene-prediction parameters and was not applied directly as coordinate-specific evidence on the cultivar consensus sequences. AUGUSTUS was run with soft-masking enabled. Predicted protein sequences were searched against the AGIS1.0 reference proteome using DIAMOND blastp v2.1.16 (Buchfink et al. 2021) in sensitive mode, with an E-value threshold of 1e-5, a minimum identity percentage of 50%, query coverage of 0%, and a maximum target sequence of 1. The highest-scoring retained hit for each query was used for annotation. Hits were classified as informative or as uncharacterized/hypothetical according to an annotation-based rule, while queries without a retained hit were reported separately. Gene annotations were converted from GFF to BED format using the GFF-to-BED converter in Galaxy Europe (https://usegalaxy.eu) (The Galaxy Community, 2026) and checked against the predicted peptide identifiers. BEDTools v2.31.1 (Quinlan and Hall, 2010) was used to generate provisional strand-aware 2-kb upstream intervals for coordinate inspection (using ‘bedtools slopbed’). These provisional intervals were used only as quality-control inputs. The promoter sequences used for the final AGIS1.0-anchored comparisons were reconstructed from complete gene coordinates using the dedicated workflow described below.

### BUSCO completeness analysis and BUSCO-based phylogenetic reconstruction

BUSCO v5.8.0 (Simão et al. 2015; Manni et al. 2021) was used for completeness analysis, including the assessment of complete single and duplicate orthologs, as well as fragmented and missing orthologs. BUSCO v5.8.0 was run in protein mode using the embryophyta_odb10 dataset to assess complete single-copy, complete duplicated, fragmented, and missing orthologs. The BUSCO orthologs across cultivars were compared and displayed as a percentage bar chart. For phylogenomic reconstruction, BUSCO loci classified as complete single-copy in all eight datasets were retained. Sequence headers were standardized, and loci were regrouped using rename_busco_headers.py and group_buscos_by_id.py from BUSCO Supermatrix Helper (see Data Availability). Individual loci were aligned using MAFFT v7.525 (Katoh 2002, 2005), producing 1565 peptide loci. Alignments were concatenated using concatenate_busco_alignments.py, yielding final matrices across cultivars. Maximum-likelihood trees were reconstructed using IQ-TREE v2.4.0 (Nguyen et al. 2014) with ModelFinder Plus (-m MFP), 1,000 ultrafast-bootstrap replicates (-B 1000) (Minh et al. 2013), and 1,000 SH-aLRT replicates (--alrt 1000) (Guindon et al. 2010). The selected substitution models were [model names]. The generated trees based on nucleotides and peptides were compared for analysis. The tree was visualized using MEGA 11 software as presented in Figure 3.

### Orthogroup inference and comparative predicted proteome analysis

Predicted protein sequences generated with AUGUSTUS for the seven Indonesian rice cultivars were analyzed together with the predicted proteome of the AGIS1.0 reference genome using OrthoFinder v2.5.5 (Emms and Kelly 2019). Inclusion of AGIS1.0 provided a common reference for interpreting orthogroup composition and species relationships across the cultivar datasets. OrthoFinder was run using its standard workflow to assign proteins to orthogroups and generate orthogroup-level statistics, per-species assignment statistics, pairwise species-overlap summaries, and a species tree. The topology of the OrthoFinder species tree was subsequently compared with that obtained from the concatenated BUSCO phylogenomic analysis. The tree was also visualized using MEGA 11 software.

For comparative gene-content analysis, orthogroup presence was determined from OrthoFinder output using Python scripts from Shared Orthogroup Analyzer (see Data Availability). Orthogroup classes used only the seven Indonesian cultivars, with AGIS1.0 as a reference. Orthogroups in all seven cultivars were ‘core’, in six ‘semi-core’, in 2-5 ‘accessory’, and in one ‘private’. Those absent in all cultivars but present in AGIS1.0 were ‘reference-only’. Because genomes were reference-guided, these categories reflect predicted orthogroup patterns, not validated gene presence-absence. Orthogroup-sharing patterns were visualized with an UpSet plot from the binary matrix. Exact orthogroup intersections were calculated from the binary presence-absence matrix for the seven Indonesian cultivars, excluding AGIS1.0 from the cultivar-based class definitions. The main UpSet plot displayed the core intersection shared by all seven cultivars, the ten largest intersections with occupancy between two and six cultivars, and all seven cultivar-private intersections. Complete intersection counts were retained in the accompanying machine-readable output.

### Trait-related candidate-gene recovery and predicted protein comparison

A panel of 40 AGIS1.0 loci or gene-family entries associated with amino-acid and nitrogen metabolism, grain pigmentation, and starch properties was compiled from published gene names, locus identifiers, protein accessions, functional descriptions, and genomic coordinates (Table S1). Nine chalcone synthase loci were treated separately to avoid collapsing adjacent paralogues.

Custom Python scripts were generated to integrate the target list with the AGIS1.0 and cultivar AUGUSTUS peptide FASTA files, gene-coordinate BED files, OrthoFinder orthogroup assignments, and DIAMOND tabular results (see Data Availability). The candidate-recovery script identified the orthogroup containing each AGIS1.0 target, retrieved its cultivar members, and ranked them using chromosome correspondence, distance from the reference locus, and within-chromosome gene order. Candidates on the corresponding chromosome and within 1 Mb of the AGIS1.0 locus were prioritized. For tandem or fragmented loci, genomic position and model proximity were used to distinguish paralogues and select local model clusters. The script produced a cultivar-by-target matrix containing selected model identifiers, coordinates, orthogroup and DIAMOND evidence, recovery status, model multiplicity, predicted peptide length, and annotation-quality flags (Tables S2 and S6). Model quality was classified from the number of selected models, the ratio of their combined predicted peptide length to the corresponding AGIS1.0 target length, and the proportion of unknown amino-acid residues. Peptides containing more than 5% unknown (X) residues were classified as low confidence. For single-model loci, length ratios below 0.75 were classified as truncated or partial, ratios from 0.75 to below 0.85 as short or potentially partial, ratios from 0.85 to 1.20 as full-length candidates, and ratios above 1.20 as overextended or potentially chimeric. Multi-model loci were classified as split but incomplete below 0.75, likely split from 0.75 to 1.20, and split or overextended above 1.20.

A second script retrieved the selected peptide sequences and compared each cultivar candidate with its corresponding AGIS1.0 model. Model quality was classified from model number, peptide-length agreement, and the proportion of unknown residues. Direct residue-wise comparison was restricted to single-model reference and cultivar peptides of equal length. Direct positional comparison was restricted to single-model AGIS1.0 and cultivar peptides of equal length. For these compatible pairs, the script calculated positional identity and reported amino-acid differences relative to the predicted AGIS1.0 peptide. Split, fragmented, or length-discordant models were reported as gene-model or predicted-length differences and were not subjected to residue-wise comparison (Table S3). DIAMOND results provided independent protein-level support but were not used alone to establish locus identity. Failure to recover a defensible model was not interpreted as biological gene deletion without read-backed validation.

### Reference-anchored promoter extraction, alignment, and quality control

Candidate loci were reconstructed from the selected gene-model table. Multiple models assigned to the same target were combined into one locus only when they occurred on the same contig and strand and were separated by no more than 5 kb. Loci spanning different contigs or strands, or containing selected models separated by more than 5 kb, were excluded when a single defensible promoter boundary could not be established. For each retained locus, a strand-aware 2-kb upstream interval was extracted from the corresponding genome FASTA using the complete gene coordinates. For plus-strand genes, the interval [gene start − 2,000, gene start) was extracted; for minus-strand genes, [gene end, gene end + 2,000) was extracted and reverse-complemented. Intervals extending beyond a contig boundary were truncated and flagged.

Each cultivar promoter was compared independently with its corresponding AGIS1.0 promoter using a script for the promoter analysis (see Data Availability). To evaluate whether differences in predicted gene boundaries displaced the apparent promoter interval, exact 31-mers were sampled every 10 bp from the AGIS1.0 promoter. Only 31-mers occurring once in the reference promoter and once within a local query-genome window were retained. Candidate anchor positions were clustered within 120 bp, and an anchor was considered strong when supported by at least eight hits representing at least 4% of eligible unique reference 31-mers. When a strong anchor displaced the query interval by at least 50 bp from its predicted boundary, the syntenically anchored 2-kb interval was used; otherwise, the predicted-boundary interval was retained. Offsets of at least 100 bp were flagged for gene-boundary review.

Promoter pairs were globally aligned using a Needleman-Wunsch implementation with a match score of +2, mismatch score of −3, and linear gap penalty of −4. Alignment-derived SNVs and contiguous insertion or deletion events were recorded relative to the AGIS1.0 interval. These differences represent pairwise alignment calls and not independently validated read-level variants. Comparisons were flagged when either gene model was not classified as full-length, an interval was shorter than 2 kb, syntenic-anchor support was weak, the anchor offset was at least 100 bp, the selected interval overlapped the predicted query gene boundary by more than 50 bp, global identity was below 95%, or either interval contained ambiguous N bases. A comparison was classified as passing quality control only when none of these conditions applied.

A curated catalogue of exact IUPAC cis-element words was scanned on both DNA strands. Motif counts and apparent gain or loss events were evaluated relative to aligned orthologous positions. Because short or degenerate motif words may occur by chance, these results were treated as exploratory candidates and not as evidence of transcription-factor binding or altered gene expression.

## Results

### PacBio HiFi sequencing generated high-coverage datasets for seven rice cultivars

PacBio HiFi sequencing of the seven Indonesian rice cultivars generated a combined 95.40 Gb from a total of 19,144,502 reads (Table 1). Sequencing yield per cultivar ranged from 10.77 Gb in Cempo Ireng (CI) to 16.04 Gb in Gogo Jak (GJ), with 2.40-3.07 million HiFi reads obtained per cultivar. Mean read lengths ranged from 4,494 bp in CI to 5,667 bp in GJ, while read N50 values ranged from 5,438 bp in Inpago (IN) to 7,015 bp in GJ. Based on the AGIS1.0 reference-genome size, the datasets represented estimated coverages of 27.92×-41.58×, with a mean of 35.34× across cultivars. GJ produced the greatest sequencing yield, estimated coverage, mean read length, and read N50, while all seven cultivars generated broadly comparable read depth for the subsequent reference-guided analyses.

**Table 1.** PacBio HiFi sequencing summary.

| Cultivar ID | Total Bases (bp) | Total HiFi Reads | Mean Read Length (bp) | N50 (bp) | Coverage Estimated to AGIS1.0 (×) |
| --- | --- | --- | --- | --- | --- |
| IN | 11360897664 | 2527728 | 4494.51 | 5438 | 29.46 |
| CI | 10767697040 | 2395765 | 4494.47 | 5449 | 27.92 |
| GJ | 16037029531 | 2829656 | 5667.48 | 7015 | 41.58 |
| KM | 14544461159 | 3071709 | 4734.97 | 5790 | 37.71 |
| MPE | 13846408142 | 2626895 | 5271.02 | 6436 | 35.90 |
| BSM | 14352594026 | 2627334 | 5462.80 | 6887 | 37.21 |
| PP | 14495222423 | 3065415 | 4728.63 | 5841 | 37.58 |

### Reference-guided consensus genomes were constructed for all cultivars

Mapping of the PacBio HiFi reads to AGIS1.0 produced reported mapping rates of 100% for all seven cultivars, with mean mapping-quality values ranging from 51.17 in BSM to 54.79 in KM (Table 2). Mean sequencing depth ranged from 26.74× in CI to 40.76× in GJ, with an overall mean of 34.02×. The resulting consensus genomes ranged from 387.93 Mb in KM to 390.54 Mb in BSM, representing a narrow size difference of 2.61 Mb across the seven cultivars. GJ retained the greatest mapped depth, consistent with its higher sequencing yield, whereas the remaining cultivars showed depths of 26.74× −36.95×. These datasets therefore provided broadly comparable reference-guided genomic representations for downstream completeness, repeat, annotation, phylogenomic, and orthogroup analyses. Because positions without a retained alternative call remain in the AGIS1.0 state, the resulting sequences represent cultivar-specific, reference-guided consensus resources rather than independently assembled genomes. The narrow range of consensus-sequence sizes consequently reflects, in part, their shared reference framework.

**Table 2.**
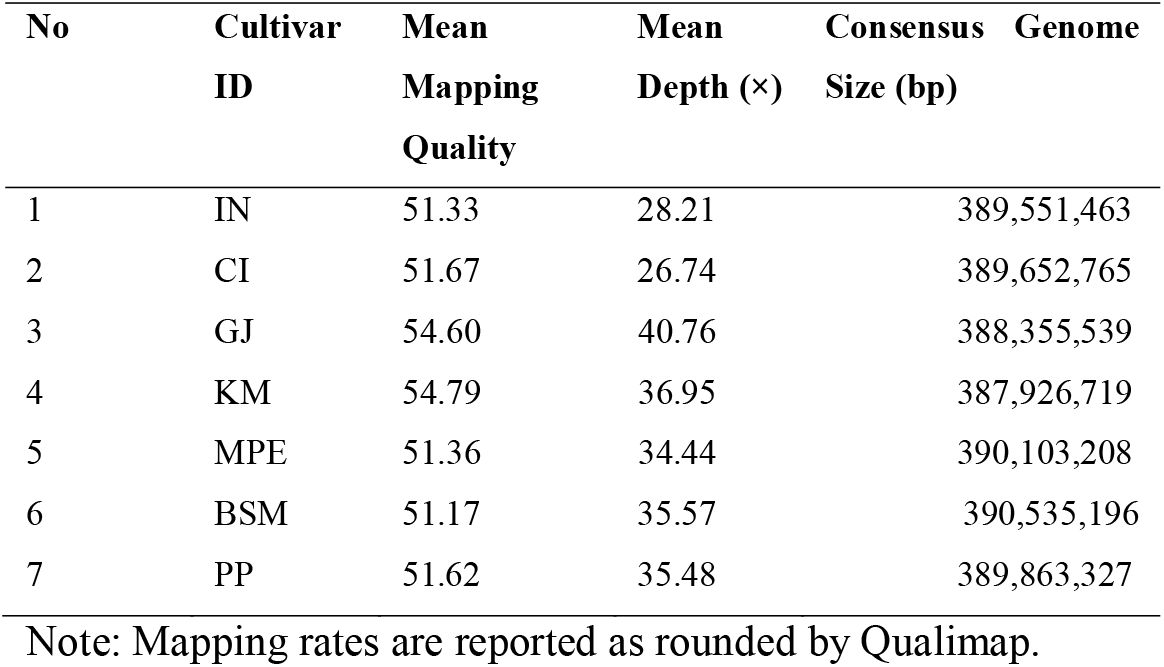
Read-mapping and reference-guided consensus-genome statistics for seven Indonesian rice cultivars.

### Genome completeness and repeat landscapes were broadly conserved among cultivars

BUSCO analysis showed highly similar gene-space completeness across the seven reference-guided consensus genomes (Fig. 1A). Complete BUSCOs accounted for approximately 98.3-98.5% of the assessed orthologs, comprising approximately 96.0-96.4% complete single-copy and 2.2-2.5% complete duplicated BUSCOs. Fragmented and missing BUSCOs each represented approximately 0.5-0.7% and 1.0-1.1%, respectively. AGIS1.0 showed a slightly higher complete BUSCO proportion of approximately 98.6%, but its overall profile remained closely comparable to those of the seven cultivars. These values indicate that the reference-guided consensus sequences retained nearly complete representation of the conserved gene space already present within the AGIS1.0 coordinate framework. They should not be interpreted as independent estimates of de novo assembly completeness.

**Fig 1.**
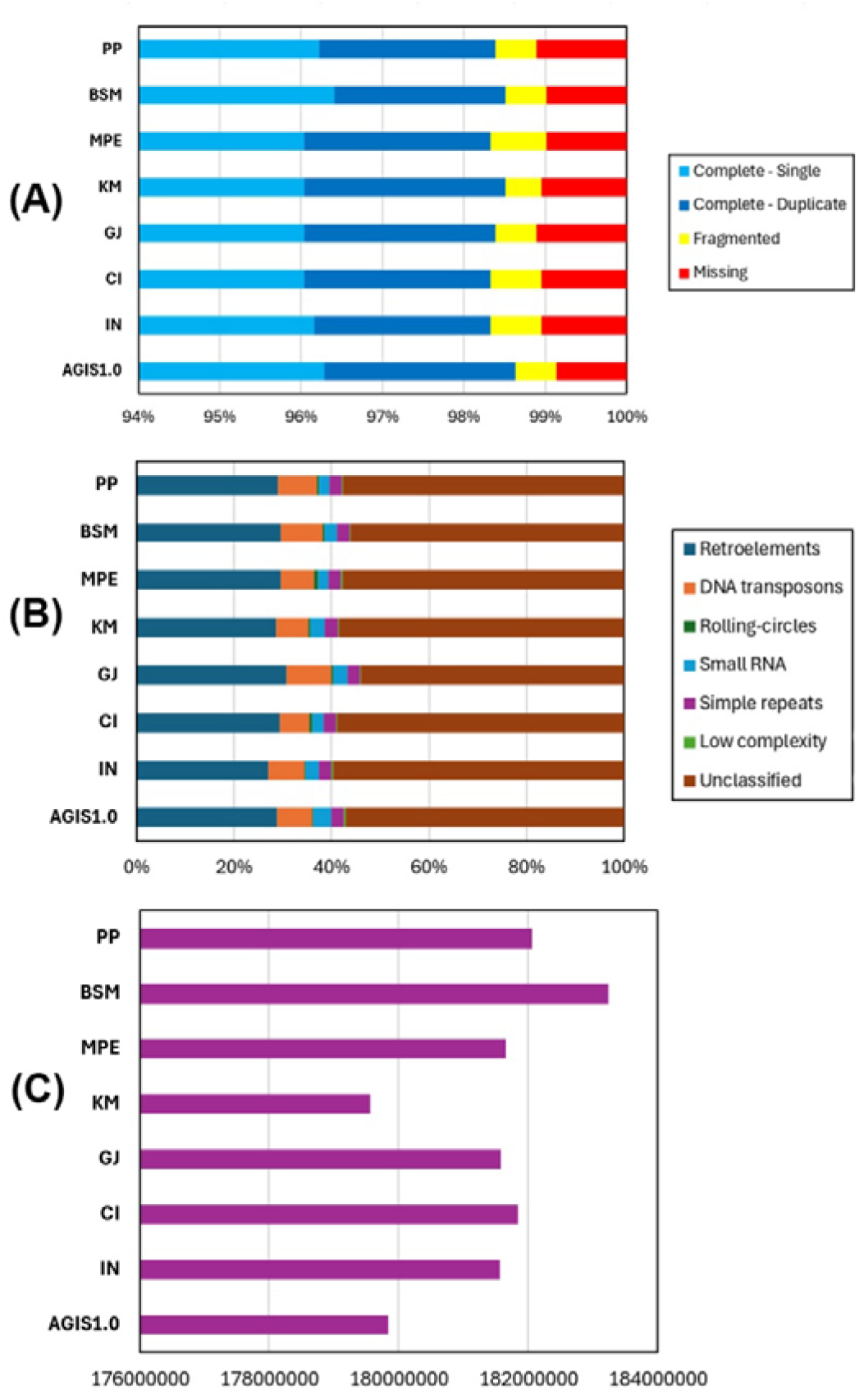
BUSCO gene-space completeness (A), relative composition of RepeatMasker classes (B), and total interspersed-repeat length (C) across AGIS1.0 and the seven reference-guided cultivar consensus sequences.

Repeat profiles were likewise broadly similar among datasets (Fig. 1B-C). Unclassified repeats formed the largest component of the displayed repeat categories, followed by retroelements and DNA transposons, while rolling-circle elements, small-RNA-associated repeats, simple repeats, and low-complexity sequences constituted smaller fractions. The total length of interspersed repeats varied within a narrow range of approximately 179.6-183.3 Mb: KM had the lowest value and BSM the highest, while AGIS1.0 contained approximately 179.8 Mb. Within the reference-guided consensus framework, no dataset showed a pronounced difference in total interspersed-repeat length. This comparison does not evaluate non-reference insertions or the genomic positions of individual repeat elements.

### Structural and functional annotation produced comparable gene sets across cultivars

AUGUSTUS predicted 39,178-40,150 protein-coding gene models across the seven reference-guided cultivar consensus sequences, compared with 39,737 models in AGIS1.0 (Table 3; Fig. 2). Among the cultivars, 21,784-22,035 models received informative DIAMOND annotations, while 8,435-8,670 matched proteins described as uncharacterized or hypothetical. The remaining 8,955-9,445 models had no DIAMOND hit under the applied search criteria. KM had the largest predicted gene set and the greatest number of informatively annotated models, whereas GJ had the smallest predicted gene set and the fewest models without a DIAMOND hit. Across AGIS1.0 and the seven cultivars, the mean counts were 21,896.13 informatively annotated models, 8,523.13 models with uncharacterized or hypothetical hits, and 9,216.38 models without a DIAMOND hit. The narrow ranges demonstrate consistent output from the shared reference-guided and gene-prediction workflow, rather than independently establishing equivalent biological gene content among the cultivars.

**Fig. 2.**
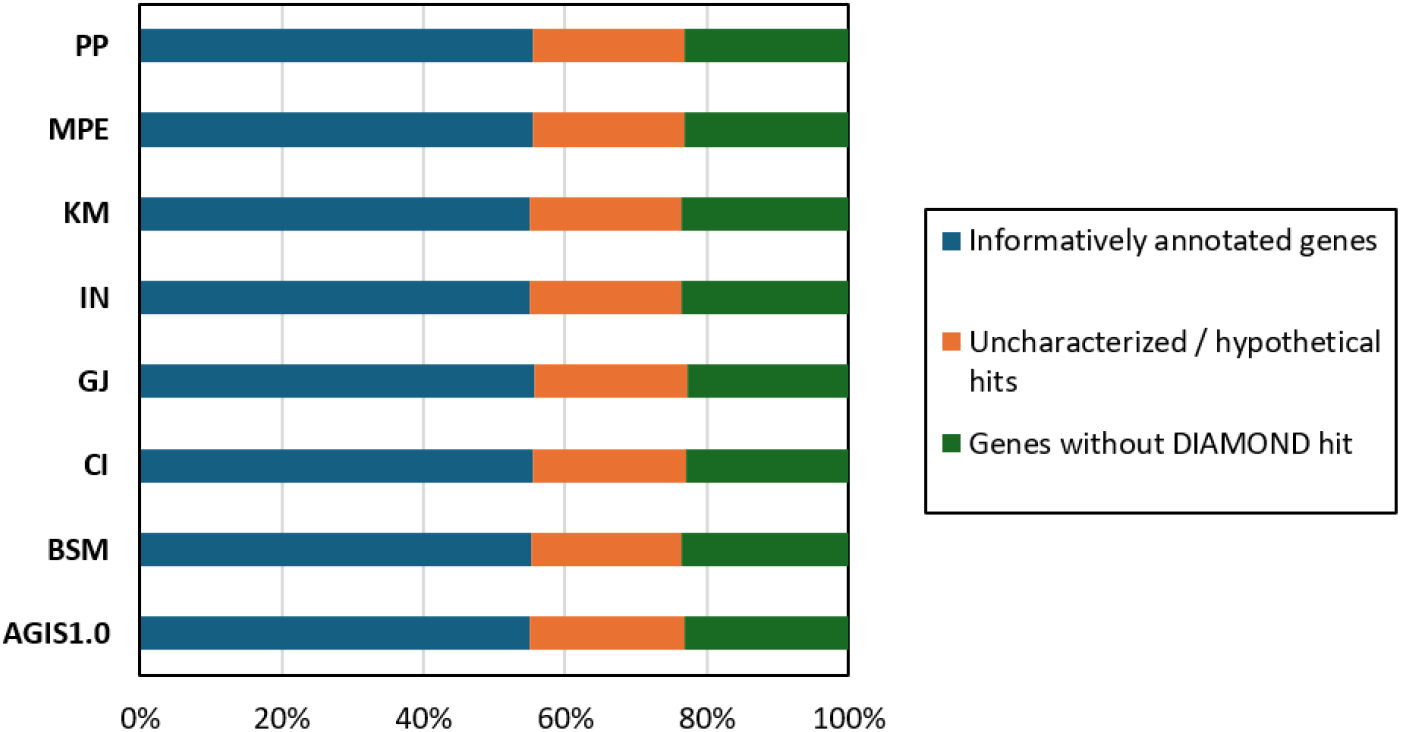
Relative distribution of predicted gene models with informative DIAMOND hits, uncharacterized or hypothetical protein hits, or no retained DIAMOND hit across AGIS1.0 and the seven cultivar consensus sequences.

**Table 3.**
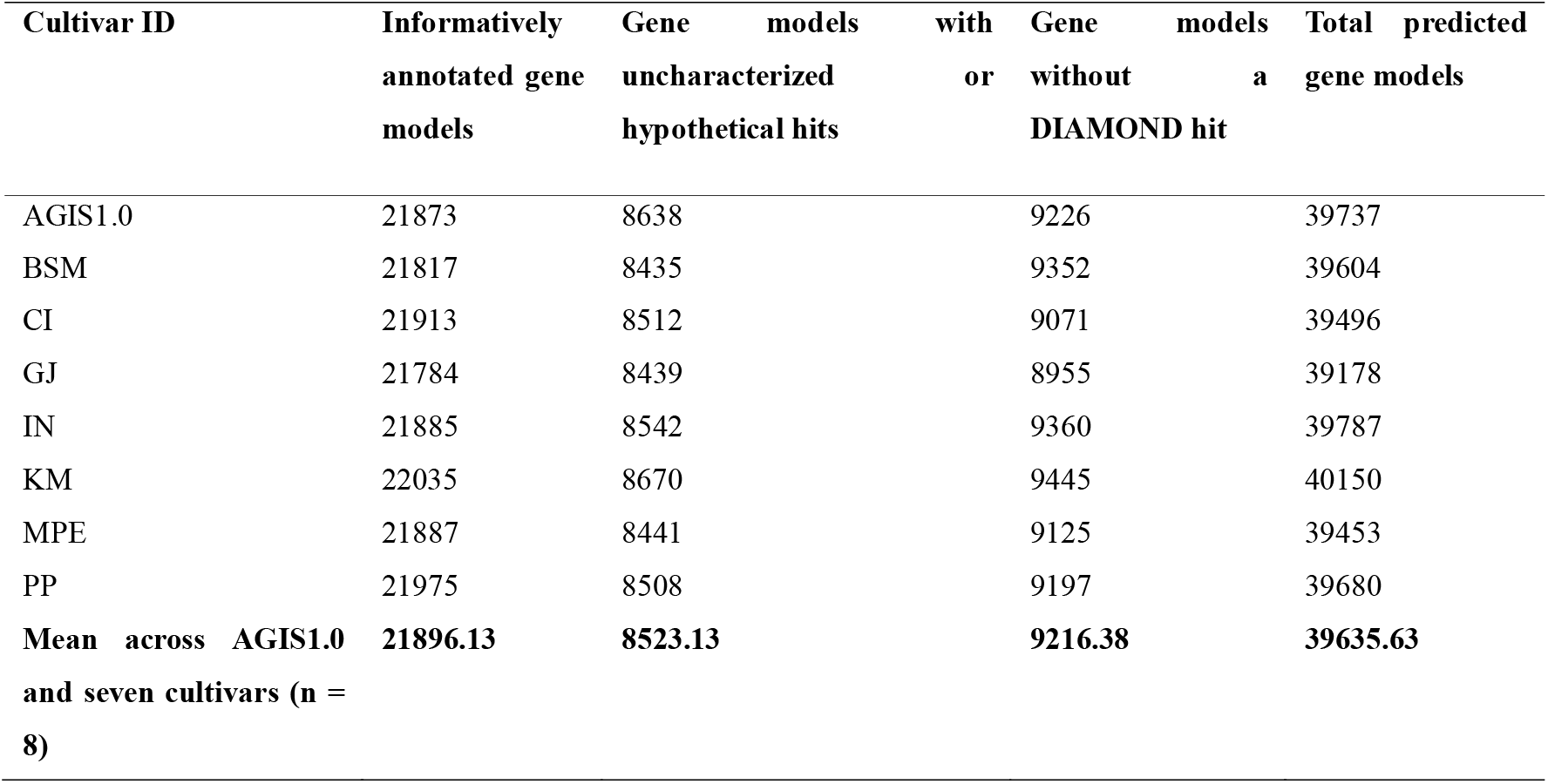
Summary of AUGUSTUS gene models classified according to DIAMOND annotation outcome.

### BUSCO-based phylogeny revealed conserved gene-space relationships among rice cultivars

The maximum-likelihood tree reconstructed from concatenated BUSCO orthologs recovered short branches among all eight datasets (Fig. 3). AGIS1.0 and GJ formed the closest pair, with KM placed adjacent to them, whereas BSM and CI formed a second close pair within the remaining group. CI and MPE, the two pigmented cultivars highlighted in the figure, did not form an exclusive pair. Because the cultivar sequences were constructed against AGIS1.0 and the analysis sampled conserved BUSCO loci, this topology represents similarity among reference-guided consensus sequences rather than cultivar ancestry or population structure.

**Fig. 3.**
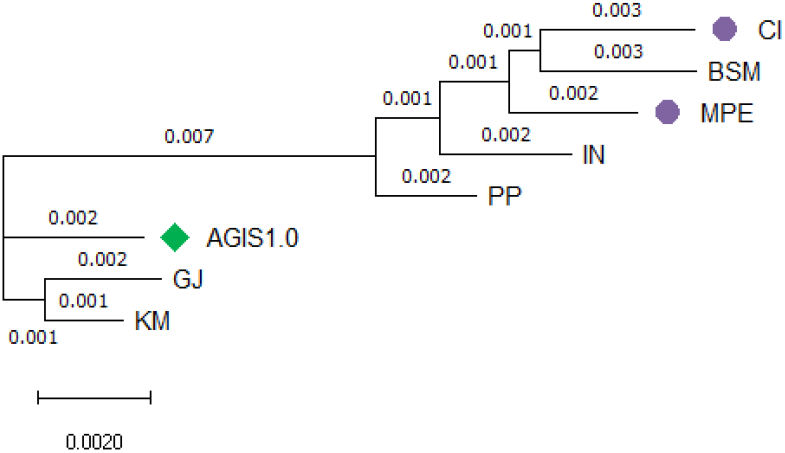
Maximum-likelihood phylogeny reconstructed from concatenated BUSCO orthologs. Node labels indicate SH-aLRT and ultrafast-bootstrap support, and the scale bar represents substitutions per site. The green diamond indicates AGIS1.0, and purple circles indicate the two pigmented cultivars.

### Orthogroup inference revealed a large conserved predicted core and limited private representation

OrthoFinder assigned 314,321 of 317,085 predicted proteins (99.1%) to 40,737 orthogroups (Fig. 4). Individual cultivars were represented in 34,303-34,909 orthogroups, with GJ having the lowest and KM the highest count. Considering only the seven Indonesian cultivars, 27,514 orthogroups were classified as core, 2,692 as semi-core, 9,720 as accessory, and 792 as private. An additional 19 orthogroups were detected only in AGIS1.0. Thus, 67.5% of all inferred orthogroups were represented across all seven cultivars, indicating a large conserved component of the predicted gene space.

**Fig. 4.**
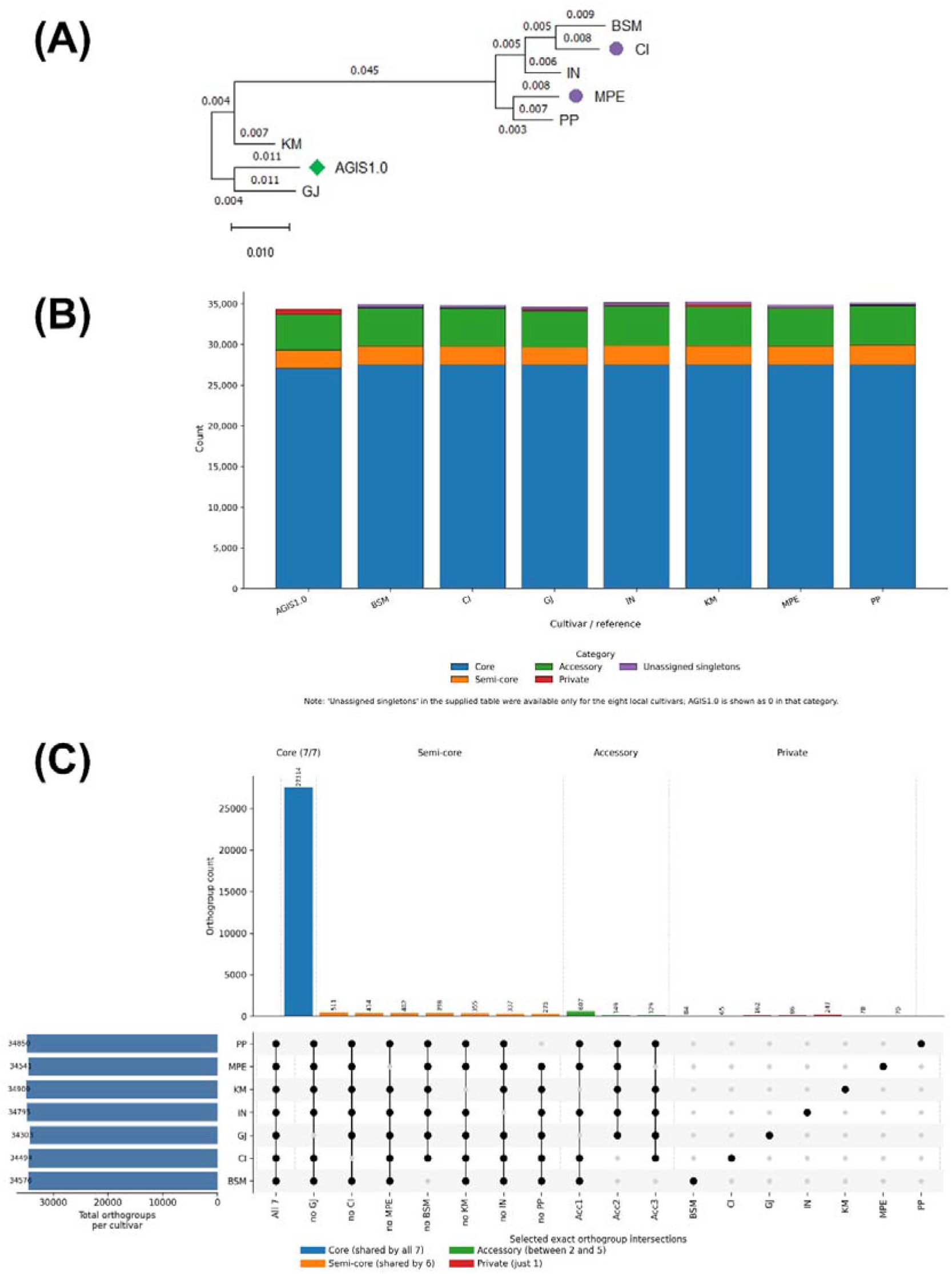
OrthoFinder species tree (A), distribution of cultivar-represented orthogroup classes (B), and selected exact orthogroup intersections among the seven Indonesian cultivars (C). AGIS1.0 was included in OrthoFinder but excluded from the seven-cultivar core, semi-core, accessory, and private definitions. Panel C displays the core intersection, the ten largest intersections shared by two to six cultivars, and all cultivar-private intersections. The green diamond indicates AGIS1.0, and purple circles indicate the two pigmented cultivars.

Private orthogroup representation was limited but varied among cultivars, ranging from 65 orthogroups in CI to 247 in KM. The remaining counts were 70 in PP, 78 in MPE, 84 in BSM, 86 in IN, and 162 in GJ. The largest exact intersection comprised the 27,514 core orthogroups shared across all seven cultivars, whereas the selected semi-core and accessory intersections were substantially smaller (Fig. 4B–C). Because orthogroup assignment depends on the predicted protein models, private, accessory, and reference-only classifications should be interpreted as differences in predicted orthogroup recovery rather than confirmed gene gain, loss, or cultivar-specific presence–absence variation.

The OrthoFinder species tree recovered the same principal relationships observed in the BUSCO-based phylogeny (Fig. 4A). AGIS1.0 and GJ formed the closest pair, with KM placed next to them, while BSM and CI formed a second close pair. MPE and PP also grouped, with IN placed adjacent to this pair. The repeated recovery of the AGIS1.0-GJ-KM and BSM-CI groupings indicates that both methods captured a similar conserved gene-space signal. It does not provide independent confirmation of cultivar genealogy because the two analyses used related predicted coding-sequence information derived from the same reference-guided resources.

### Targeted trait-related genes were broadly recovered and revealed focused sequence candidates

Forty target loci or family entries associated with amino-acid transport and nitrogen metabolism, pigmentation, and starch biosynthesis or properties were screened using AGIS1.0-anchored orthogroup membership, chromosome-collinear coordinates, predicted protein-model quality, and DIAMOND similarity searches. Across the seven Indonesian cultivars, 278 of 280 target-by-cultivar combinations were recovered, of which 277 were DIAMOND-supported. All 217 non-CHS target-by-cultivar combinations were recovered and supported by DIAMOND. The exceptions were confined to the CHS family: CHS_L3 and CHS_L9 were not recovered in CI by the current orthogroup-plus-synteny screen, whereas the short 96-aa CHS_L7 candidate in KM lacked a DIAMOND hit. Systematic abnormalities affecting OsGS2/GLN2, CHI, DFR, and SBE1/BEI were classified as gene-model quality issues rather than evidence of biological gene gain, fusion, or loss (Fig. 5A; Tables S1, S2, and S6).

**Fig. 5.**
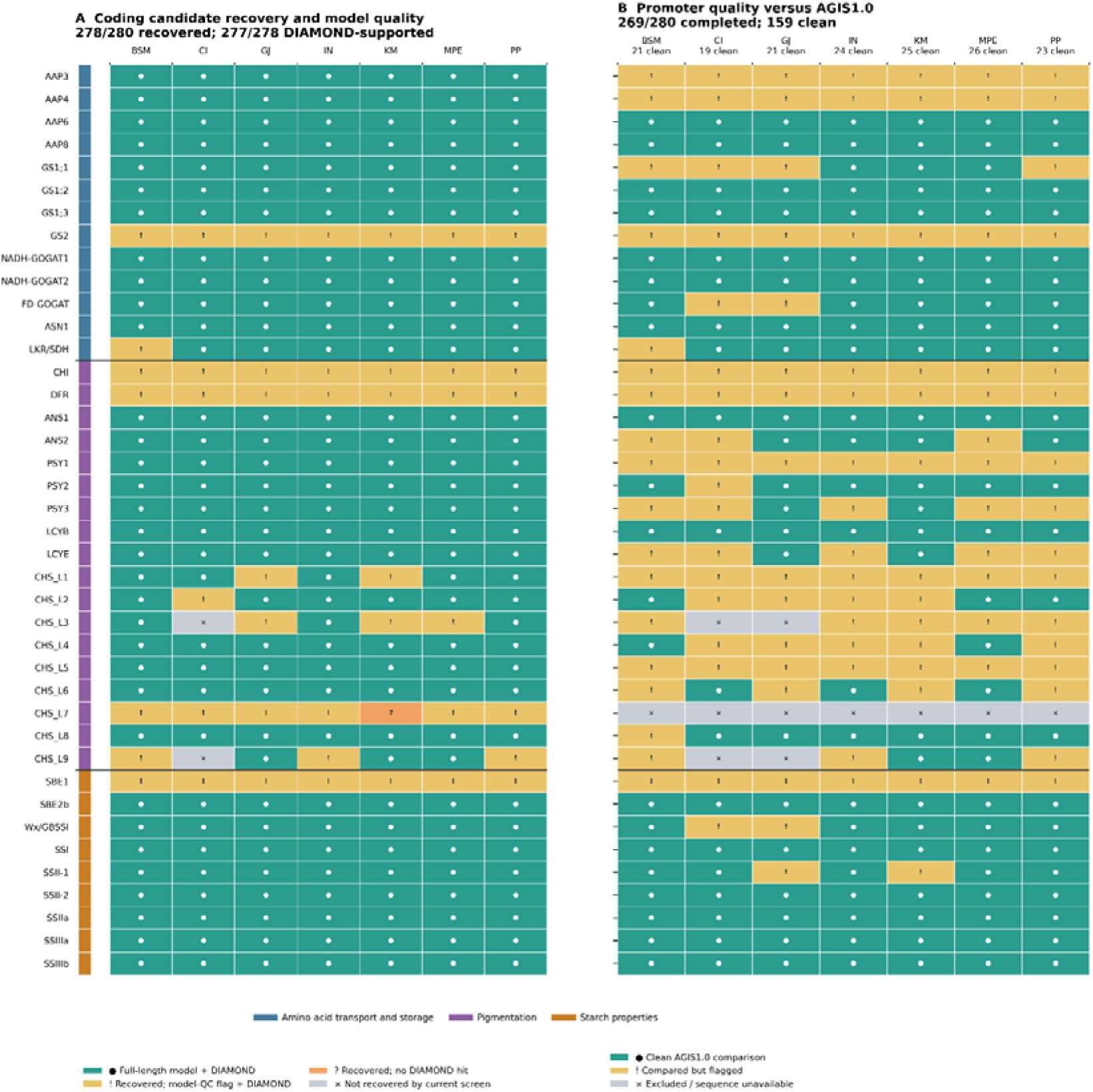
Candidate-gene recovery and predicted model quality (A) and AGIS1.0-anchored promoter-comparison quality control (B) across seven Indonesian rice cultivars. Of 280 target-by-cultivar combinations, 278 were recovered, and 277 were supported by DIAMOND. Of 280 scheduled promoter comparisons, 159 passed, 110 were flagged, and 11 were excluded. Trait groups are indicated by coloured rows.

Among structurally comparable, equal-length predicted protein models, CI, IN, and MPE shared a 13-residue sequence pattern relative to AGIS1.0, whereas BSM and PP shared a smaller five-residue pattern. Recurrent positional amino-acid differences were also detected in starch-related proteins. BSM, CI, IN, MPE, and PP showed L94V and H196R in SBE2b, whereas GJ and KM matched AGIS1.0 at these positions. M730V in SSIIa/ALK was detected in BSM, GJ, KM, MPE, and PP, with MPE additionally showing D88E and S597G; the length-discordant CI and IN models require further curation. GJ, IN, KM, and MPE showed Y224S in Wx/GBSSI, whereas PP showed D166G, and the extended CI model remained unsuitable for direct positional comparison. BSM, CI, and PP shared five positional differences in SSI, whereas MPE showed K438E. OsAAP6/qPC1 showed a distinct predicted protein-model length pattern, with 466-aa models in BSM, MPE, and PP compared with 478-aa models in AGIS1.0, CI, GJ, IN, and KM. These comparisons prioritized ANS1, SBE2b, SSIIa/ALK, Wx/GBSSI, OsAAP6/qPC1, and SSI for read-backed and functional validation (Table 4; Fig. S1; Table S3).

**Table 4.** Principal trait-related candidate genes prioritized from predicted protein comparison and AGIS1.0-anchored promoter analysis across seven Indonesian rice cultivars.

| Candidate | Trait context | Predicted protein evidence | Promoter comparison and QC | Interpretation and required validation |
| --- | --- | --- | --- | --- |
| ANS1 | Pigmentation | CI, IN, and MPE shared a 13-residue sequence pattern; BSM and PP shared a smaller pattern | All seven comparisons passed; 99.30-99.95% identity | Strongest integrated candidate, validate nucleotide sequence, expression, enzyme function, and grain pigment composition |
| *SBE2b* | Starch branching | L94V and H196R in BSM, CI, IN, MPE, and PP; GJ and KM matched AGIS1.0 | All seven passed; 97.77-100% identity | High-priority starch candidate, motif implications remain exploratory |
| *SSIIa/ALK* | Starch properties | M730V in BSM, GJ, KM, MPE, and PP; additional differences in MPE; CI and IN models were length-discordant | All seven passed; 97.09-99.90% identity | Curate CI and IN gene models and test gelatinization and starch phenotypes |
| *Wx/GBSSI* | Amylose synthesis | Y224S in GJ, IN, KM, and MPE; D166G in PP; CI model was extended | Five passed at 97.62-99.75%; CI and GJ flagged at 49.67% and 50.07% | Important predicted protein candidate, flagged promoter intervals require boundary and structural review |
| *OsAAP6/qPC1* | Grain protein and nitrogen | 466-aa models in BSM, MPE, and PP versus 478-aa models in the other datasets | All seven passed; CI, GJ, IN, and KM matched AGIS1.0, while BSM, MPE, and PP retained clean differences | Confirm exon structure and transcript sequence and test grain protein and amino-acid content |
| *SSI* | Starch synthesis | Five shared positional differences in BSM, CI, and PP; K438E in MPE | All seven passed; 98.26-100% identity | Secondary starch candidate requiring starch chain-length and physicochemical phenotyping |

Each selected 2-kb promoter interval was aligned independently to its corresponding AGIS1.0 reference interval. Of 280 scheduled comparisons, 269 were completed: 159 passed the sequence-, model-, and boundary-quality criteria, 110 were completed but flagged, and 11 were excluded because a defensible selected promoter sequence was unavailable (Fig. 5B; Table S4). The excluded comparisons comprised all seven CHS_L7 comparisons, for which no defensible AGIS1.0 promoter was available, together with the CI and GJ comparisons for CHS_L3 and CHS_L9. These exclusions do not demonstrate gene or promoter deletion. The 269 completed comparisons contained 19,604 alignment-derived SNVs and 8,850 contiguous indel events, whereas the 159 clean comparisons contained 1,811 SNVs and 803 indel events. Thus, although flagged comparisons represented 40.9% of completed comparisons, they contained 90.8% of the SNVs and 90.9% of the indel events, indicating that most apparent divergence was concentrated in loci with gene-model, promoter-boundary, synteny, or structural concerns.

*ANS1*, *SBE2b*, *SSIIa/ALK*, *OsAAP6/qPC1*, and *SSI* passed promoter quality control in all seven AGIS1.0-versus-cultivar comparisons. Their minimum promoter identities were 99.30%, 97.77%, 97.09%, 96.16%, and 98.26%, respectively. Five of the seven *Wx/GBSSI* comparisons also passed quality control, with identities of 97.62-99.75%, whereas the selected CI and GJ promoter intervals showed 49.67% and 50.07% identity, respectively. These two comparisons were therefore classified as promoter-boundary or structural-review cases rather than credible regulatory-sequence differences (Fig. S2). For *OsAAP6*, the selected CI, GJ, IN, and KM promoter intervals were identical to AGIS1.0, whereas BSM, MPE, and PP retained clean sequence differences. The predicted protein-model length difference, rather than a shared promoter-divergence pattern, therefore represented the principal *OsAAP6* sequence signal.

Reference-anchored comparisons also resolved which cultivars generated the apparent sequence divergence at problematic loci. The selected *OsAAP4* promoter matched AGIS1.0 in GJ and KM but showed only approximately 76.5-76.9% identity in BSM, CI, IN, MPE, and PP. Because the AGIS1.0 locus itself contains a short or potentially partial gene model, all *OsAAP4* promoter comparisons were retained as validation cases. *LCYE* was nearly identical to AGIS1.0 in GJ and KM but produced low-identity selected intervals in BSM, CI, IN, MPE, and the selected PP interval was shifted by 942 bp relative to the predicted reference boundary. *OsGS1;1* passed the available quality checks in IN, KM, and MPE but showed low identity or substantial promoter-boundary displacement in BSM, CI, GJ, and PP. These loci were therefore retained as QC and validation cases and were excluded from direct regulatory interpretation until their gene models, local synteny, and promoter boundaries can be confirmed using mapped reads or local assembly (Fig. S3; Table S6).

## Discussion

This study provides a reference-guided comparison of seven Indonesian rice cultivars and reveals a consistent overall pattern: broadly similar reference-represented genome and predicted gene-space profiles accompanied by a smaller set of cultivar-dependent orthogroup, predicted protein-sequence, and promoter differences. The high sequencing depths, near-complete BUSCO profiles, comparable repeat totals, similar predicted model counts, and recovery of 278 of 280 targeted loci demonstrate consistent recovery of conserved AGIS1.0-represented gene space through a shared analytical workflow. These results do not establish equivalent complete genome content outside the reference framework. Nevertheless, the orthogroup analysis and focused candidate screening identified testable differences potentially relevant to grain pigmentation, starch properties, and protein content. These resources expand the genomic characterization of Indonesian rice germplasm, which contains substantial diversity associated with subspecies, cultivation system, geography, and local selection (Thomson et al. 2009).

The PacBio HiFi datasets provided approximately 28-42× estimated coverage per cultivar, supporting reliable read mapping and small-variant consensus construction. HiFi sequencing combines long reads with high per-base accuracy and is therefore well suited to detecting sequence variants and resolving complex genomic regions (Wenger et al. 2019). AGIS1.0 was also an appropriate coordinate framework because it represents a complete telomere-to-telomere assembly of the Nipponbare genome and resolves regions absent or misassembled in earlier rice references (Shang et al. 2023). However, the reported 100% mapping rates and narrow range of consensus-genome sizes should not be interpreted as evidence that the seven cultivars are nearly identical at the whole-genome level. The consensus sequences were generated through alignment to AGIS1.0, and positions without supported alternative calls retain the reference state. Consequently, their similar sizes, BUSCO profiles, and repeat landscapes partly reflect the shared reference framework as well as genuine conservation.

This reference dependence is particularly important when interpreting sequences that are absent from, structurally different from, or poorly collinear with Nipponbare. Large rice surveys have identified extensive SNP, indel, structural-variant, gene copy-number, and presence–absence diversity that cannot be represented completely by a single linear reference (Wang et al. 2018; Qin et al. 2021). Long-read sequencing of 111 rice accessions recovered substantially more non-reference sequence than short-read pan-genomes, with repetitive sequence forming a major component of the newly recovered regions (Zhang et al. 2022). The present consensus genomes should therefore be considered high-coverage, cultivar-specific representations of sequence variation relative to AGIS1.0, rather than independent de novo assemblies or a complete Indonesian rice pan-genome.

BUSCO completeness remained approximately 98.3-98.5% across the cultivar consensus genomes, and annotation produced 39,178-40,150 predicted protein-coding models per cultivar. These narrow ranges demonstrate the internal consistency of the common processing and annotation workflow. The repeat results similarly showed no pronounced genome-wide expansion or contraction among the consensus sequences. However, similar total repeat contents do not exclude extensive variation in the positions of individual transposable elements. More than 50,000 retrotransposon insertion polymorphisms have been identified across the 3,000 Rice Genomes collection, illustrating that positional variation may remain extensive even when total repeat abundance is similar (Carpentier et al. 2019). Cultivar-specific de novo assemblies and read-backed transposable-element insertion analysis would therefore be required to determine whether the Indonesian cultivars possess distinctive repeat or structural-variant landscapes.

The annotation results also demonstrate why predicted gene-model counts alone are insufficient for inferring biological gene-content differences. The systematic split, truncated, or overextended models observed for OsGS2/GLN2, CHI, DFR, and SBE1/BEI occurred across multiple datasets and were more consistent with annotation behaviour than with repeated independent gene disruption. Likewise, the 8,955–9,445 models without DIAMOND hits may include lineage-specific or poorly characterized proteins, but may also include incomplete or erroneous predictions. Transcript evidence, full-length cDNA or Iso-Seq data, conserved-domain analysis, and local read-backed reconstruction would be required to distinguish genuine genes from gene-model artefacts. Accordingly, the annotation set is most appropriately treated as a standardized comparative resource and a source of testable candidates rather than as a final catalogue of functional genes.

OrthoFinder assigned 99.1% of predicted proteins to orthogroups and identified 27,514 core orthogroups represented in all seven cultivars. This large core is consistent with the close evolutionary relationships expected among cultivated rice accessions. The repeated AGIS1.0-GJ-KM and BSM-CI groupings indicate that the BUSCO and OrthoFinder analyses captured a similar conserved gene-space signal. Because both analyses used coding information derived from the same reference-guided datasets, their agreement should not be regarded as independent evidence of cultivar genealogy. However, neither topology should be interpreted as a definitive cultivar genealogy. The seven genomes were constructed against the same Nipponbare reference, the trees were based predominantly on conserved coding regions, and the sampling was too limited for population-genetic inference. The finding that the two pigmented cultivars, CI and MPE, did not form an exclusive group further indicates that grain pigmentation represents a locus- and pathway-dependent phenotype rather than a marker of overall genomic relatedness.

The accessory and private orthogroup categories provide an initial view of variable predicted gene representation, but they require similar caution. Private counts ranged from 65 orthogroups in CI to 247 in KM, while only 19 orthogroups were reference-only. Some of these patterns may represent genuine sequence divergence, structural variation, or gene presence–absence variation, all of which are widespread in rice pan-genomes (Qin et al. 2021). Others may result from gene-model fragmentation, failure to predict a protein, or sequence divergence affecting orthogroup assignment. Read-depth inspection, nucleotide-level synteny, local assembly, and comparison against de novo cultivar genomes will be necessary before individual orthogroups can be described as genuine gene gains, losses, or cultivar-specific sequences.

The targeted analysis showed that the examined amino-acid metabolism, pigmentation, and starch-related gene families were overwhelmingly conserved. The two unrecovered CI CHS entries and the short KM CHS_L7 model should therefore be interpreted as locus-recovery or annotation exceptions, rather than demonstrated deletions. Within the recovered loci, ANS1 was the most prominent pigmentation-related predicted protein-sequence candidate. CI, IN, and MPE shared a 13-residue pattern relative to AGIS1.0, whereas BSM and PP shared a smaller five-residue pattern. ANS1 participates in anthocyanin biosynthesis, but pigment accumulation depends on coordinated expression of structural genes and transcriptional regulators. In black rice, for example, ectopic expression caused by structural rearrangement of the Kala4 promoter can activate the broader anthocyanin pathway (Oikawa et al. 2015). The presence of the same ANS1 pattern in IN as well as the pigmented cultivars CI and MPE further suggests that this protein pattern alone is unlikely to explain grain colour. Its relevance should be tested through nucleotide confirmation, developing-grain expression analysis, enzyme-level assessment, and anthocyanin profiling.

The recurrent variation observed in SBE2b, SSIIa/ALK, Wx/GBSSI, and SSI is biologically plausible because these enzymes have established roles in rice endosperm starch synthesis. Loss or reduction of SBE2b activity alters amylopectin branching and can increase apparent amylose content (Nishi et al. 2001), while ALK/SSIIa is a major determinant of starch gelatinization temperature (Gao et al. 2003). SSI contributes particularly to the synthesis of short and intermediate amylopectin chains (Fujita et al. 2006), and natural Wx alleles produce substantial differences in amylose biosynthesis and eating quality (Zhang et al. 2019). The SBE2b L94V/H196R pattern, SSIIa M730V pattern, Wx Y224S pattern, and SSI substitutions identified here are therefore suitable targets for follow-up. Nevertheless, these specific substitutions have not been demonstrated to alter enzyme activity, starch structure, cooking quality, or glycaemic properties. Functional interpretation will require validated nucleotide haplotypes together with measurements of amylose content, amylopectin chain-length distribution, gelatinization properties, and starch digestibility.

OsAAP6/qPC1 represents a similarly plausible grain-protein candidate. OsAAP6 has been experimentally demonstrated to regulate amino-acid uptake and distribution and to influence rice grain protein content (Peng et al. 2014). In the present analysis, BSM, MPE, and PP were represented by 466-aa models rather than the 478-aa models found in AGIS1.0 and the other cultivars. In contrast, their promoter comparisons did not reveal a single shared divergent pattern, and four cultivar promoter intervals were identical to AGIS1.0. The predicted predicted protein-model length difference is therefore the principal OsAAP6 signal, but it could represent either a genuine coding or splice difference or a systematic gene-model discrepancy. Read-backed exon reconstruction, full-length transcript sequencing, and grain protein and amino-acid measurements are necessary before this pattern can be interpreted functionally.

Reference-anchored promoter comparison improved the biological clarity of the analysis by evaluating each cultivar independently against AGIS1.0 and by separating clean comparisons from problematic intervals. Although flagged comparisons constituted 40.9% of the completed set, they contained 90.8% of all alignment-derived SNVs and 90.9% of all contiguous indel events. This concentration demonstrates that unfiltered estimates of promoter divergence can be dominated by gene-model, boundary, synteny, or structural problems. In contrast, ANS1, SBE2b, SSIIa/ALK, OsAAP6, and SSI passed the available quality criteria in all seven cultivars and generally retained high promoter identity. The approximately 50% identities of the selected CI and GJ Wx intervals, together with recurrent anomalies involving OsAAP4, LCYE, and OsGS1;1, are more appropriately treated as boundary or structural-review cases than as exceptionally divergent regulatory sequences. This distinction is important because promoter-boundary definition can strongly affect both sequence comparison and motif discovery (Ksouri et al. 2021).

Even within clean intervals, sequence differences do not establish regulatory effects. A short motif-word gain or loss may occur without changing transcription-factor binding, while genuine expression differences can depend on chromatin context, distal regulatory elements, interacting transcription factors, or developmental stage. Conversely, experimentally validated rice traits show that specific promoter changes can have major phenotypic consequences, as demonstrated for Kala4 and OsAAP6 (Oikawa et al. 2015; Peng et al. 2014). The promoter results should consequently be regarded as a prioritization framework. Expression profiling in developing grain, reporter assays, transcription-factor binding experiments, and targeted promoter editing would be required to establish regulatory function.

Several limitations define the boundary of the present conclusions. The study included seven cultivars without a population-level sampling design, formal association testing, transcriptomic evidence, or matched biochemical phenotypes. The consensus sequences were unphased and reference-guided, the annotations were produced computationally, and proximal 2-kb intervals cannot capture the complete regulatory landscape. A logical next stage would combine read-backed local assembly and structural-variant calling at the prioritized loci with de novo HiFi assembly of each cultivar. These assemblies could then support a graph-based Indonesian rice pan-genome, more reliable gene presence–absence and transposable-element analyses, and haplotype-resolved candidate comparisons. Developing-grain transcriptomics and targeted phenotyping would subsequently allow the ANS1, starch-related, and OsAAP6 patterns to be tested against pigment composition, grain protein content, and starch physicochemical properties.

Overall, this study establishes a reference-guided genomic baseline for seven Indonesian rice cultivars and demonstrates that focused candidate differences can be prioritized within a largely conserved predicted gene space. Its principal contribution is not the demonstration of cultivar-specific causal alleles, but the creation of standardized genomic resources and a ranked collection of predicted protein, promoter, annotation, and structural hypotheses. These resources provide a foundation for read-backed locus reconstruction, transcriptomic and biochemical investigation, and the incorporation of locally maintained Indonesian rice diversity into future conservation and breeding programmes.

## Conclusion

This study establishes a reference-guided comparative genomic baseline for seven Indonesian rice cultivars. The shared workflow recovered highly consistent AGIS1.0-represented gene space and prioritized predicted protein and promoter-sequence candidates associated with pigmentation, grain protein content, and starch properties. ANS1, SBE2b, SSIIa/ALK, Wx/GBSSI, OsAAP6/qPC1, and SSI emerged as the strongest candidates for subsequent investigation. Systematic quality control further showed that most apparent promoter divergence occurred in flagged intervals affected by gene-model, boundary, synteny, or possible structural problems. The resulting candidates should therefore be regarded as testable hypotheses rather than confirmed functional alleles. Because the sequences were constructed against AGIS1.0, future de novo and haplotype-resolved assemblies, developing-grain transcriptomics, biochemical phenotyping, and functional assays will be required to establish causal genotype–phenotype relationships and support the use of this local rice diversity in conservation and breeding.

## Acknowledgment

We thank members of the Pigmented Rice Research Group, Research Center for Biotechnology, Universitas Gadjah Mada, for their support and scientific discussions. We also acknowledge Yayasan Satriabudi Dharma Setia (YSDS) and the Integrated Genomic Facility (IGF), Faculty of Biology, Universitas Gadjah Mada, for supporting PacBio whole-genome sequencing.

## Declarations AI Usage

OpenAI ChatGPT was used to assist with computational workflow development, quality-control design, figure and table generation, and manuscript drafting and language editing. Using author-supplied AUGUSTUS annotations, BED files, OrthoFinder results, DIAMOND results, promoter sequences, and reference-annotation outputs, the tool helped develop and refine Python workflows for candidate-gene integration, model-quality classification, predicted protein comparison, AGIS1.0-anchored promoter comparison, variant summarization, and quality-control reporting. The AI did not generate sequencing data or provide experimental validation. All scripts, parameters, candidate assignments, numerical outputs, figures, interpretations, and manuscript text were inspected and approved by the authors, who retain responsibility for the study design, accuracy, biological interpretation, and final content. Validated scripts are available through the repositories listed in the Data Availability statement. Grammarly was used for grammatical and language checking.

## Supplementary Data

**Figure S1.** Coding variation in six principal candidate genes across seven Indonesian rice cultivars.

**Figure S2.** Reference-anchored *Wx/GBSSI* promoter comparisons across seven Indonesian rice cultivars.

**Figure S3.** Recurrent promoter-boundary and structural-review loci across the seven AGIS1.0-versus-cultivar comparisons

**Table S1**. Candidate-gene definitions and AGIS1.0 reference loci.

**Table S2**. Coding-candidate recovery, DIAMOND support, and gene-model QC.

**Table S3**. Peptide substitutions and model-length differences relative to AGIS1.0.

**Table S4**. All 280 scheduled AGIS1.0-versus-cultivar promoter comparisons.

**Table S5**. Exploratory exact motif-word gain/loss events.

**Table S6**. Annotation, promoter-boundary, and structural-variation validation flags.

## Data Availability

PacBio HiFi reads generated for the seven cultivars are available through the NCBI Sequence Read Archive under BioProject PRJNA1493835. Custom analysis scripts are available from the BUSCO Supermatrix Helper repository (https://github.com/adhitwicaksono/busco-supermatrix-helper), Shared Orthogroup Analyzer repository (https://github.com/adhitwicaksono/SharedOrthogroupAnalyzer), Rice Trait Candidate Gene Analyzer repository (https://github.com/adhitwicaksono/rice-trait-candidate-gene-analyzer), and promoter-analysis repository (https://github.com/adhitwicaksono/promoter-analyzers).

## References

Buchfink B, Reuter K, Drost HG. 2021. Sensitive protein alignments at tree-of-life scale using DIAMOND. Nature Methods 18:366–368. 10.1038/s41592-021-01101-x

Carpentier MC, Manfroi E, Wei FJ, et al. 2019. Retrotranspositional landscape of Asian rice revealed by 3000 genomes. Nature Communications 10:24. 10.1038/s41467-018-07974-5

Danecek P, Bonfield JK, Liddle J, et al. 2021. Twelve years of SAMtools and BCFtools. GigaScience 10:giab008. 10.1093/gigascience/giab008

Emms, D.M. and Kelly, S., 2019. OrthoFinder: phylogenetic orthology inference for comparative genomics. Genome Biology, 20(1), p.238. 10.1186/s13059-019-1832-y

Flynn JM, Hubley R, Goubert C, et al. 2020. RepeatModeler2 for automated genomic discovery of transposable element families. Proceedings of the National Academy of Sciences 117:9451–9457. 10.1073/pnas.1921046117

Fujita N, Yoshida M, Asakura N, et al. 2006. Function and characterization of starch synthase I using mutants in rice. Plant Physiology 140:1070–1084. 10.1104/pp.105.071845

Gao Z, Zeng D, Cui X, et al. 2003. Map-based cloning of the *ALK* gene, which controls the gelatinization temperature of rice. Science in China Series C: Life Sciences 46:661–668. 10.1360/03yc0099

Guindon, et al. (2010) New Algorithms and Methods to Estimate Maximum-Likelihood Phylogenies: Assessing the Performance of PhyML 3.0. Systematic Biology 59, 3, 307–321, 10.1093/sysbio/syq010

Hoang DT, Chernomor O, von Haeseler A, Minh BQ, Vinh LS. 2018. UFBoot2: improving the ultrafast bootstrap approximation. Molecular Biology and Evolution 35:518–522. 10.1093/molbev/msx281

Hoff, K.J. and Stanke, M., 2019. Predicting genes in single genomes with AUGUSTUS. Current Protocols in Bioinformatics, 65(1), e57. 10.1002/cpbi.57

Katoh, K. (2002). MAFFT: a novel method for rapid multiple sequence alignment based on fast Fourier transform. Nucleic Acids Research, 30(14), 3059–3066. 10.1093/nar/gkf436

Katoh, K. (2005). MAFFT version 5: improvement in accuracy of multiple sequence alignment. Nucleic Acids Research, 33(2), 511–518. 10.1093/nar/gki198

Ksouri N, Castro-Mondragón JA, Montardit-Tarda F, van Helden J, Contreras-Moreira B, Gogorcena Y. 2021. Tuning promoter boundaries improves regulatory motif discovery in nonmodel plants: the peach example. Plant Physiology 185:1242–1258. 10.1093/plphys/kiaa091

Li, H., Handsaker, B., Wysoker, A., Fennell, T., Ruan, J., Homer, N., Marth, G., Abecasis, G., & and, R. D. (2009). The Sequence Alignment/Map format and SAMtools. Bioinformatics, 25(16), 2078– 2079. 10.1093/bioinformatics/btp352

Manni, M., Berkeley, M.R., Seppey, M., Simão, F.A. and Zdobnov, E.M., 2021. BUSCO update: novel and streamlined workflows along with broader and deeper phylogenetic coverage for scoring of eukaryotic, prokaryotic, and viral genomes. Molecular Biology and Evolution, 38(10), pp.4647–4654. 10.1093/molbev/msab199

Minh BQ, Schmidt HA, Chernomor O, et al. 2020. IQ-TREE 2: new models and efficient methods for phylogenetic inference in the genomic era. Molecular Biology and Evolution 37:1530–1534. 10.1093/molbev/msaa015

Nishi A, Nakamura Y, Tanaka N, Satoh H. 2001. Biochemical and genetic analysis of the effects of amylose-extender mutation in rice endosperm. Plant Physiology 127:459–472. 10.1104/pp.010127

Nguyen, L.-T., Schmidt, H. A., von Haeseler, A., & Minh, B. Q. (2014). IQ-TREE: A Fast and Effective Stochastic Algorithm for Estimating Maximum-Likelihood Phylogenies. Molecular Biology and Evolution, 32(1), 268–274. 10.1093/molbev/msu300

Oikawa T, Maeda H, Oguchi T, et al. 2015. The birth of a black rice gene and its local spread by introgression. The Plant Cell 27:2401–2414. 10.1105/tpc.15.00310

Okonechnikov, K., Conesa, A., & Garcıa-Alcalde, F. (2015). Qualimap 2: advanced multi-sample quality control for high-throughput sequencing data. Bioinformatics, btv566. 10.1093/bioinformatics/btv566

Peng B, Kong H, Li Y, et al. 2014. OsAAP6 functions as an important regulator of grain protein content and nutritional quality in rice. Nature Communications 5:4847. 10.1038/ncomms5847

Qin P, Lu H, Du H, et al. 2021. Pan-genome analysis of 33 genetically diverse rice accessions reveals hidden genomic variations. Cell 184:3542–3558.e16. 10.1016/j.cell.2021.04.046

Quinlan, A. R., & Hall, I. M. (2010). BEDTools: a flexible suite of utilities for comparing genomic features. Bioinformatics, 26(6), 841–842. 10.1093/bioinformatics/btq033

Salsinha YCF, Sebastian A, Sutiyanti E, Purwestri YA, Indradewa D, Rachmawati D. 2022. The relationship between morpho-physiological changes and expression of transcription factors in NTT local rice cultivars as a response to drought stress. Indonesian Journal of Biotechnology 27(1). 10.22146/ijbiotech.65728

Sebastian A, Nugroho IC, Putra HSD, et al. 2022. Identification and characterization of drought-tolerant local pigmented rice from Indonesia. Physiology and Molecular Biology of Plants 28:1061–1075. 10.1007/s12298-022-01185-5

Sedeek K, Zuccolo A, Fornasiero A, et al. 2023. Multi-omics resources for targeted agronomic improvement of pigmented rice. Nature Food 4:366–371. 10.1038/s43016-023-00742-9

Shang L, He W, Wang T, et al. 2023. A complete assembly of the rice Nipponbare reference genome. Molecular Plant 16:1232–1236. 10.1016/j.molp.2023.08.003

Shen, W., Le, S., Li, Y., & Hu, F. (2016). SeqKit: A Cross-Platform and Ultrafast Toolkit for FASTA/Q File Manipulation. PLOS ONE, 11(10), e0163962. 10.1371/journal.pone.0163962

Simão, F. A., Waterhouse, R. M., Ioannidis, P., Kriventseva, E. V., & Zdobnov, E. M. (2015). BUSCO: assessing genome assembly and annotation completeness with single-copy orthologs. Bioinformatics, 31(19), 3210–3212. 10.1093/bioinformatics/btv351

Stanke, M., Diekhans, M., Baertsch, R., & Haussler, D. (2008). Using native and syntenically mapped cDNA alignments to improve de novo gene finding. Bioinformatics, 24(5), 637–644. 10.1093/bioinformatics/btn013

The Galaxy Community (2026). Galaxy for accessible, reproducible, and collaborative data analyses: 2026 update.” Nucleic Acids Research 54(W1): W105–W116. 10.1093/nar/gkag469

Thomson MJ, Polato NR, Prasetiyono J, et al. 2009. Genetic diversity of isolated populations of Indonesian landraces of rice (*Oryza sativa* L.) collected in East Kalimantan on the island of Borneo. Rice 2:80–92. 10.1007/s12284-009-9023-1

Wang W, Mauleon R, Hu Z, et al. 2018. Genomic variation in 3,010 diverse accessions of Asian cultivated rice. Nature 557:43–49. 10.1038/s41586-018-0063-9

Wenger AM, Peluso P, Rowell WJ, et al. 2019. Accurate circular consensus long-read sequencing improves variant detection and assembly of a human genome. Nature Biotechnology 37:1155–1162. 10.1038/s41587-019-0217-9

Zhang C, Zhu J, Chen S, et al. 2019. Wx^lv, the ancestral allele of rice Waxy gene. Molecular Plant 12:1157–1166. 10.1016/j.molp.2019.05.011

Zhang F, Xue H, Dong X, et al. 2022. Long-read sequencing of 111 rice genomes reveals significantly larger pan-genomes. Genome Research 32:853–863. 10.1101/gr.276015.121

