## Supplementary material for "Reference-guided comparative genomics of seven Indonesian rice cultivars identifies conserved gene space and trait-associated sequence candidates": Figs S1-S3

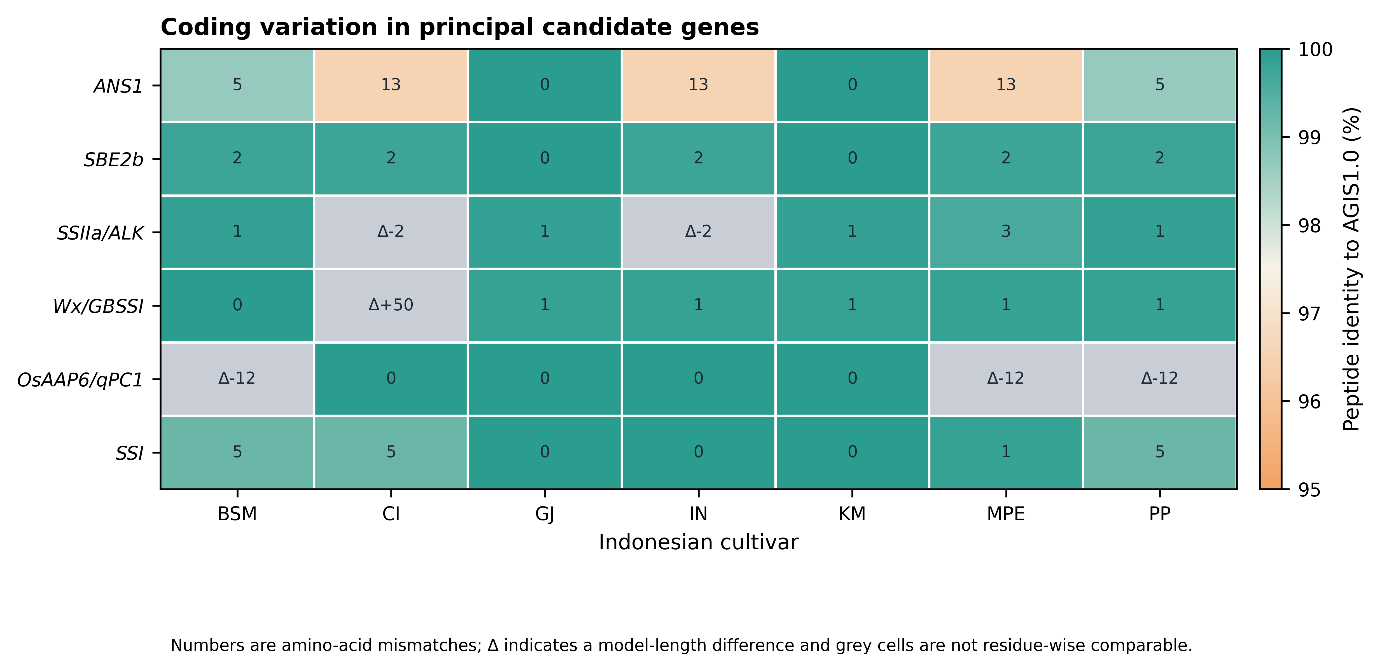

**Figure S1.** Coding variation in six principal candidate genes across seven Indonesian rice cultivars. Tile color shows peptide identity to the selected AGIS1.0 candidate where a residue-wise comparison was possible; numbers show amino-acid mismatches. Grey tiles and delta annotations indicate model-length differences or other gene-model limitations that precluded direct residue-wise comparison. These results prioritize candidates for CDS correction and nucleotide-level validation rather than establishing functional effects.

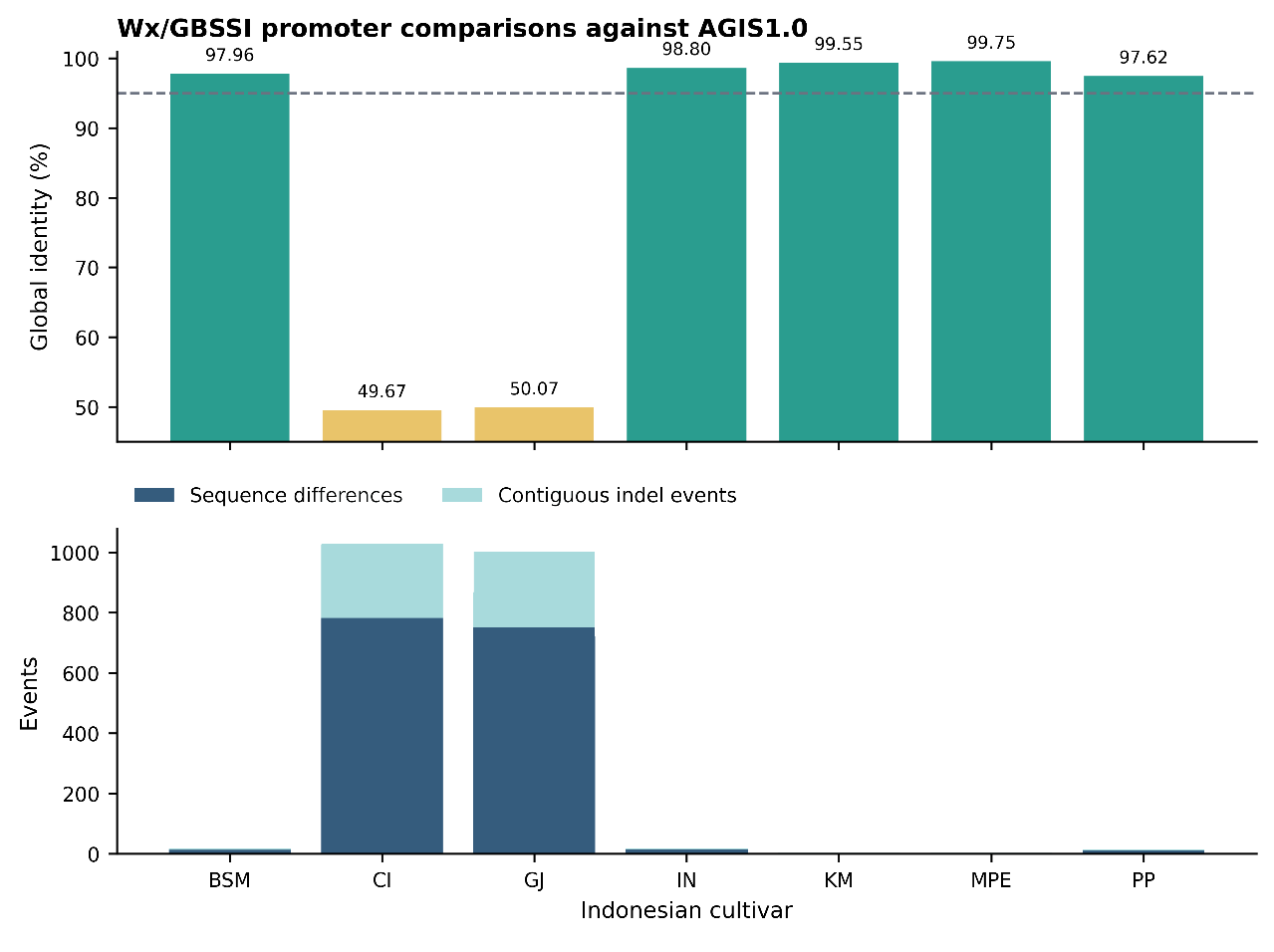

**Figure S2.** Reference-anchored *Wx/GBSSI* promoter comparisons across seven Indonesian rice cultivars. Global identity and sequence-event burden are shown for each AGIS1.0-versus-cultivar comparison. CI and GJ had markedly reduced identity (49.67% and 50.07%) and were therefore treated as promoter-boundary or structural-review cases; the other five comparisons passed the available quality checks.

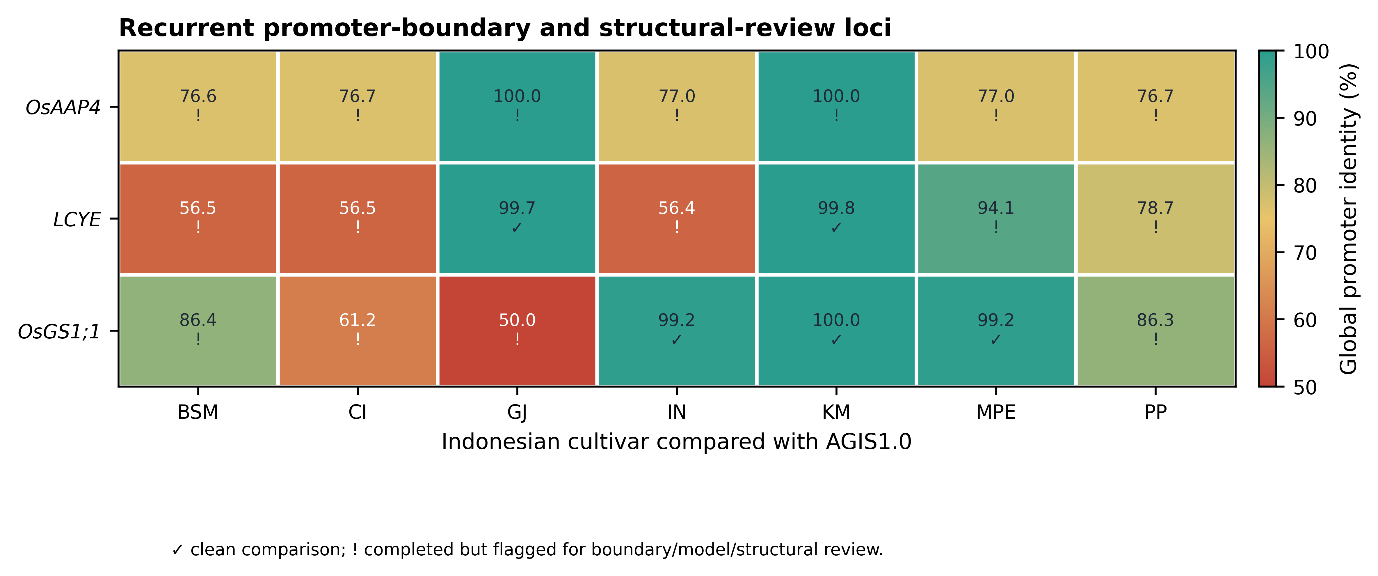

**Figure S3.** Recurrent promoter-boundary and structural-review loci across the seven AGIS1.0-versus-cultivar comparisons. Cells show global promoter identity; check marks denote clean comparisons and exclamation marks denote completed comparisons retained with sequence-, model-, or boundary-quality flags. *OsAAP4*, *LCYE*, and *OsGS1;1* were prioritized for independent local alignment, transcript-supported boundary curation, and read-backed validation before biological interpretation.
